# NASP functions in the cytoplasm to prevent histone H3 aggregation during early embryogenesis

**DOI:** 10.1101/2025.09.03.673952

**Authors:** Mohit Das, Eli Coronado-Chavez, Anusha D. Bhatt, Reyhaneh Tirgar, Amanda A. Amodeo, Jared T. Nordman

## Abstract

From their molecular birth until their incorporation into chromatin, histones are bound by specific chaperones that serve unique functions in histone trafficking, stability and chromatin deposition. The H3-specific chaperone NASP binds directly to H3 and is required to prevent degradation of soluble H3 in vivo. Where NASP functions and how NASP affects H3 dynamics and stability is unknown. Using the Drosophila early embryo as a model system to understand NASP function in vivo, we show that NASP does not directly affect H3 nuclear import or export rates. Rather, reduced H3 levels in NASP-deficient embryos indirectly affect nuclear import and the amount of H3 deposited into chromatin. Crucially, we find that cytoplasmic NASP prevents H3 aggregation in vivo and that H3 aggregation and degradation are developmentally separable events. Thus, we propose the main function of NASP in vivo is to prevent H3 aggregation, thereby indirectly protecting H3 from degradation.

## INTRODUCTION

To package the eukaryotic genome, DNA is wrapped around an octamer of histones to form nucleosomes [1–4]. Histones are not only responsible for condensing DNA to fit within the nucleus, but they also regulate access to the genetic information by controlling DNA accessibility [2, 4–6]. Histones affect nearly every aspect of chromatin metabolism and their synthesis must be carefully regulated [4–8]. Histone limitation can increase sensitivity to DNA damage and global increases in transcription [7, 9–10]. Conversely, oversupply of histones causes genomic instability, and chromosome loss in budding yeast [8, 11,12]. Excess histones can nonspecifically bind to DNA and RNA, which has the potential to impact replication, transcription and translation [8, 11, 13]. To circumvent potential histone toxicity in somatic cells, histone production peaks in S-phase when the demand is the highest and soluble (non-chromatin bound) histone pools are held to less than 1% of the total histone supply [14–19].

Embryonic development in flies, fish and frogs requires a massive and immediate supply of histones to sustain the rapid nuclear divisions typical of early embryonic development [20–22]. The early Drosophila embryo transforms from a single nucleus to 4,000-6,000 nuclei in only two hours [23, 24]. The first 13 nuclear cycles are rapid, synchronous and occur in a syncytium. During the first 13 nuclear divisions, embryogenesis is fueled by maternal deposits of proteins and RNAs, including histones [23–28]. The mid-blastula transition (MBT) occurs at nuclear cycle 14, which coincides with cell cycle slowing, cellularization, and the onset of zygotic transcription [23–24, 28–29]. In contrast to somatic cells, the vast majority of histones are soluble in the earliest stages of embryogenesis and the soluble supply slowly decreases throughout embryogenesis [20].

It is a significant challenge to measure the rate of nuclear import and export of histones in somatic cells. This is because the rate of histone nuclear import is faster than the time it takes for a fluorescent reporter such as GFP to fold and mature [17, 30]. Given histones are maternally deposited in early Drosophila embryos, nuclear import and export rates of histones have been measured and characterized in early embryos[31]. For the replication-dependent histone H3.2, the rate of nuclear import decreases every nuclear cycle due to the depletion of maternally-provided H3.2 that occurs upon each nuclear division [31, 32]. The cytoplasmic pool of H3.2 becomes limiting in nuclear cycle 13, leading to a reduction of chromatin-associated H3.2 [31]. This reduction is compensated by increased incorporation of the replication-independent H3 variant, H3.3, into chromatin starting in nuclear cycle 13 [31]. Thus, the import kinetics of H3 and its variants are concentration dependent and developmentally regulated.

To maintain the large maternal stores of histones and to suppress the potential toxicity associated with excess histones, embryos employ sophisticated mechanisms to store and regulate the availability of histones through early development. Histone chaperones are a class of proteins that associate with histones and are crucial for histone stability and histone metabolism [33, 34]. While few chaperones bind to all histone subunits, most histone chaperones are specific to H2A-H2B or H3-H4 [35–38]. In Drosophila embryos, the histone chaperone Jabba binds to H2A, H2B and H2Av and sequesters them to lipid droplets, protecting them from degradation [39]. NASP (Nuclear Autoantigenic Sperm Protein) is a histone chaperone that binds to H3-H4 reserves in both mammalian cells and Drosophila embryos and protects them from degradation [40–42]. Unlike Jabba, NASP is a maternal effect gene and embryos laid by NASP-mutant mothers have severely reduced hatching rate [39, 40, 43].

NASP binds directly to H3 through evolutionarily conserved tetratricopeptide repeat (TPR) domains and the NASP-H3 interaction has been extensively characterized [44–46]. Understanding how NASP controls H3 dynamics in vivo, however, is less well understood. Biochemical work in mammalian cells has identified a number of NASP-H3-containing complexes derived from cytosolic extracts [42, 47–48]. Subsequent work revealed that NASP localizes to the nucleoplasm and that NASP can rapidly diffuse out of the nucleus during preparation of cytosolic extracts [49]. In cells depleted of NASP, soluble pools of H3 are degraded through chaperone-mediated autophagy (CMA), which occurs in the cytoplasm [41, 50]. Taken together, it is unclear if NASP functions in the nucleus or cytoplasm to protect soluble H3 from degradation, if NASP has any direct impact on the nuclear import or export of H3 or if NASP directly protects H3 from degradation. In mammalian cells, it has been proposed that NASP could act as a nuclear receptor for H3, thereby preventing H3 from nuclear export [49]. Given the caveats of measuring the import and export dynamics of H3 in somatic cells, this model is difficult to test directly. Thus, it is still unclear how NASP functions in vivo to control the stability and trafficking of H3.

Previously, we demonstrated that the Drosophila homolog of NASP binds specifically to H3, H3.3 and H4 in both Drosophila oocytes and embryos [40]. Furthermore, H3 and H4 were both degraded in embryos devoid of NASP while H2A and H2B were unaffected [40]. Similar to somatic mammalian cells, NASP functions to protect H3 and H4 from degradation in Drosophila embryos and oocytes. Here, we leverage the Drosophila developmental system to directly measure how NASP affects the import and export dynamics of H3. We show that NASP has no direct effect on the nuclear import or export rates of H3.2 or H3.3. In embryos devoid of NASP, histone import rates are reduced but this reduction matches the expected rate based on reduced H3 protein levels. Interestingly, embryos devoid of NASP have approximately half the amount of H3 in chromatin relative to NASP proficient embryos, yet these embryos progress through nuclear cycles 10-13 with little cell cycle slowing or mitotic defects. We show that NASP is largely nucleoplasmic in Drosophila embryos but functions in the cytoplasm to stabilize H3. Finally, we show that NASP functions in the cytoplasm to prevent H3 aggregation. Thus, H3 aggregation occurs prior to degradation in vivo. Altogether, we propose that the main function of NASP in vivo is to protect H3 from aggregation and that cytoplasmic aggregates of H3 formed in the absence of NASP are targeted for degradation.

## RESULTS

### H3 nuclear import is independent of NASP

Understanding how NASP functions to control nuclear import of its direct binding partner H3 has been difficult for two reasons. First, in somatic cells, the vast majority of cellular histones is chromatin bound and less than 1% of total H3 is soluble (non-chromatin bound) [51]. This limits the amount of H3 for interaction and nuclear import studies. Second, there is a kinetic barrier in studying histone transport since the rate of nuclear histone import is faster than the folding and maturation kinetics of fluorescent proteins [52–54]. To overcome the challenges of histone tracking in somatic cells, we utilized the Drosophila early embryo as a model system. Early Drosophila embryos are stockpiled with maternal reserves of RNA and proteins and >99% of maternal deposited histones are soluble at the earliest stages of embryogenesis [20]. Thus, early embryos have large soluble pools of translated histones, serving as an excellent system to study H3 dynamics [20, 31–32].

To test if H3.2 import is dependent on NASP, we utilized the photo-switchable fluorescent reporter Dendra2 tagged to H3.2 for live imaging of early embryos laid by wild-type or *NASP-*mutant mothers. The tagged H3.2-Dendra2 is expressed from a single copy of the histone gene locus that includes all promoters, coding sequences and UTRs to ensure endogenous regulation [19]. The H3.2-Dendra2 reporter undergoes post-translational modifications in vivo and interacts with endogenous cytoplasmic binding partners [39, 55–56]. To measure the rate of H3.2 nuclear import, we performed live imaging of non-photo-switched (green) H3.2-Dendra2 in 30” intervals and calculated the total nuclear intensities across nuclear cycles 11-13 in embryos laid by wild-type or *NASP-*mutant mothers. Given the differences in histone concentration between embryos laid by wild-type or NASP mutant mothers, we measured the absolute levels of H3.2-Dendra intensities throughout nuclear cycles 11-13 (Figure 1A-B). The nuclear import rates were measured by calculating the initial slopes of nuclear H3.2-Dendra2 intensities for each nuclear cycle 11-13 [31]. We found that there is a significant decrease in the rate of H3.2-Dendra2 nuclear import for nuclear cycles 11 and 12 in embryos laid *NASP*-mutant mothers compared to embryos laid by wild-type mothers (Figure 1A-C). While the rate of nuclear cycle 13 was also reduced, it did not reach the level of significance (Figure 1A-C). Similar results were obtained when normalizing the total nuclear intensities to the maximum nuclear H3.2-Dendra2 intensity in nuclear cycle 11 immediately prior to nuclear envelope breakdown (Supplemental Figure 1A-C). NASP also binds to the non-replicative H3 variant H3.3 [31, 40]. Thus, we tested if the nuclear import of H3.3 is dependent on NASP using the same live-imaging technique for embryos tagged with H3.3-Dendra2 [31]. We observed a similar decrease in the nuclear import rate of H3.3-Dendra2 in embryos laid *NASP*-mutant mothers compared to embryos laid by wild-type mothers (Figure 1D-F; Supplemental Figure 1D-F).

**Figure 1:**
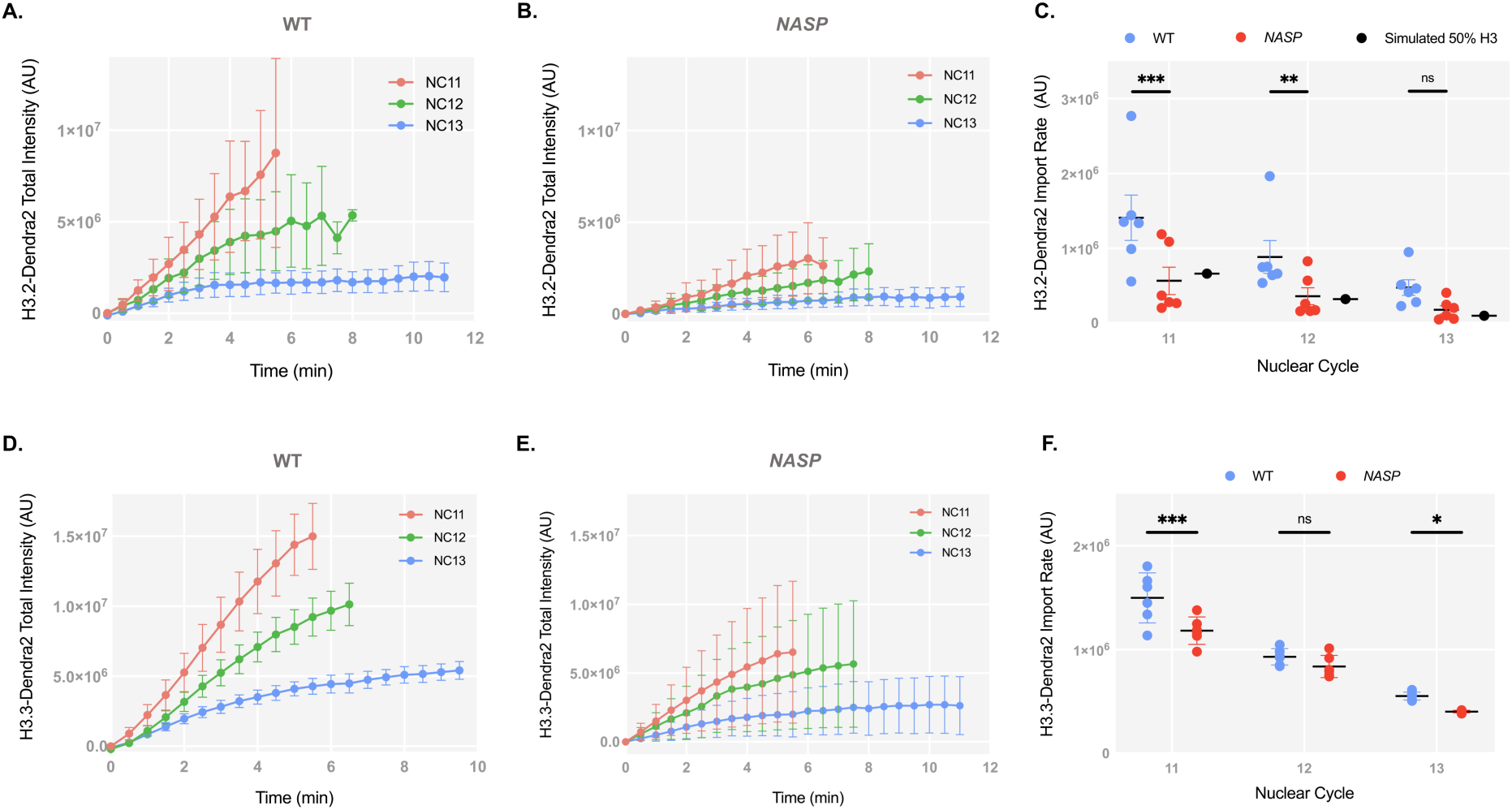
H3 nuclear import is independent of NASP. **(A)** Total intensity curves of H3.2-Dendra2 for nuclear cycles (NC) 11-13 in WT embryos. **(B)** Total intensity curves of H3.2-Dendra2 for nuclear cycles (NC) 11-13 in NASP-deficient embryos. **(C)** Nuclear import rates of H3.2-Dendra2 in WT (blue) and NASP-deficient (red) embryos. Simulated import rate for H3.2 at 50% wild-type levels (black). **(D)** Total intensity curves of H3.3-Dendra2 for nuclear cycles (NC) 11-13 in WT embryos. **(E)** Total intensity curves of H3.3-Dendra2 for nuclear cycles (NC) 11-13 in NASP-deficient embryos. **(F)** Nuclear import rates of H3.2-Dendra2 in WT (blue) and NASP-deficient (red) embryos.

The rate of H3.2 import is dependent on H3.2 concentration [32]. While the nuclear import rate of H3.2-Dendra2 was reduced in the absence of NASP, H3.2 protein levels are reduced ∼50% in embryos laid by *NASP*-mutant mothers compared to embryos laid by wild-type mothers [40]. Therefore, it is possible that the decreased H3.2 import rate in embryos laid by *NASP* mutant mothers is solely due to reduced H3.2 protein levels. To test this possibility, we used an established integrative model to simulate the rate of H3.2 nuclear import in a concentration-dependent manner [32]. We simulated H3.2 import rates with a 50% reduction of H3.2 levels to align with our experimentally measured values by mass spectrometry and western blotting [40] (Supplemental Figure 1G-H). Our experimentally derived nuclear import data aligns with simulated import rates with a 50% reduction in H3.2 concentration using either unnormalized values (Figure 1C) or values normalized to nuclear cycle 11 (Supplementary Figure 1C). Taken together, we conclude that NASP does not directly affect the nuclear import of H3 and the decrease in nuclear import rate of H3.2 in embryos laid by *NASP*-mutant mothers is solely attributed to reduced H3.2 levels.

### NASP deficient embryos have less nucleoplasmic and chromatin-associated H3

It has been suggested that NASP could function as a nuclear receptor for H3.2, thereby retaining H3.2 in the nucleus and preventing its export [48]. The measured rate of H3.2 export from the nucleus is negligible, so if there is a factor preventing H3.2 export from the nucleus, NASP is an ideal candidate [48]. We tested if NASP prevents H3.2 export by photo-converting the H3.2-Dendra2 reporter in nuclear cycle 12 and measuring the intensity of photoconverted H3.2-Dendra2 over the course of nuclear cycle 12. There was no significant change in nuclear intensity of photoconverted H3.2-Dendra2 prior to nuclear envelope breakdown for embryos laid by wild-type or *NASP*-mutant mothers, indicating that NASP does not affect export of H3.2 (Figure 2A). There is a significant reduction in H3.2-Dendra2 levels upon nuclear envelope breakdown in embryos laid by wild-type mothers due to the loss of non-chromatin bound nucleoplasmic H3.2-Dendra2 (Figure 2A) [31]. In contrast, there was only a slight reduction in H3.2-Dendra2 upon nuclear envelope breakdown in embryos laid by *NASP*-mutant mothers (Figure 2A). Similar results were observed for H3.3 (Figure 2B). Consistent with the reduced levels of total H3.2 and H3.3 in embryos laid by *NASP*-mutant mothers, these data reveal that the level of soluble nucleoplasmic H3.2 and H3.3 is reduced in *NASP*-deficient embryos. We conclude that NASP indirectly reduces both the nuclear import rate and the soluble supply of both H3.2 and H3.3 in the nucleoplasm.

**Figure 2:**
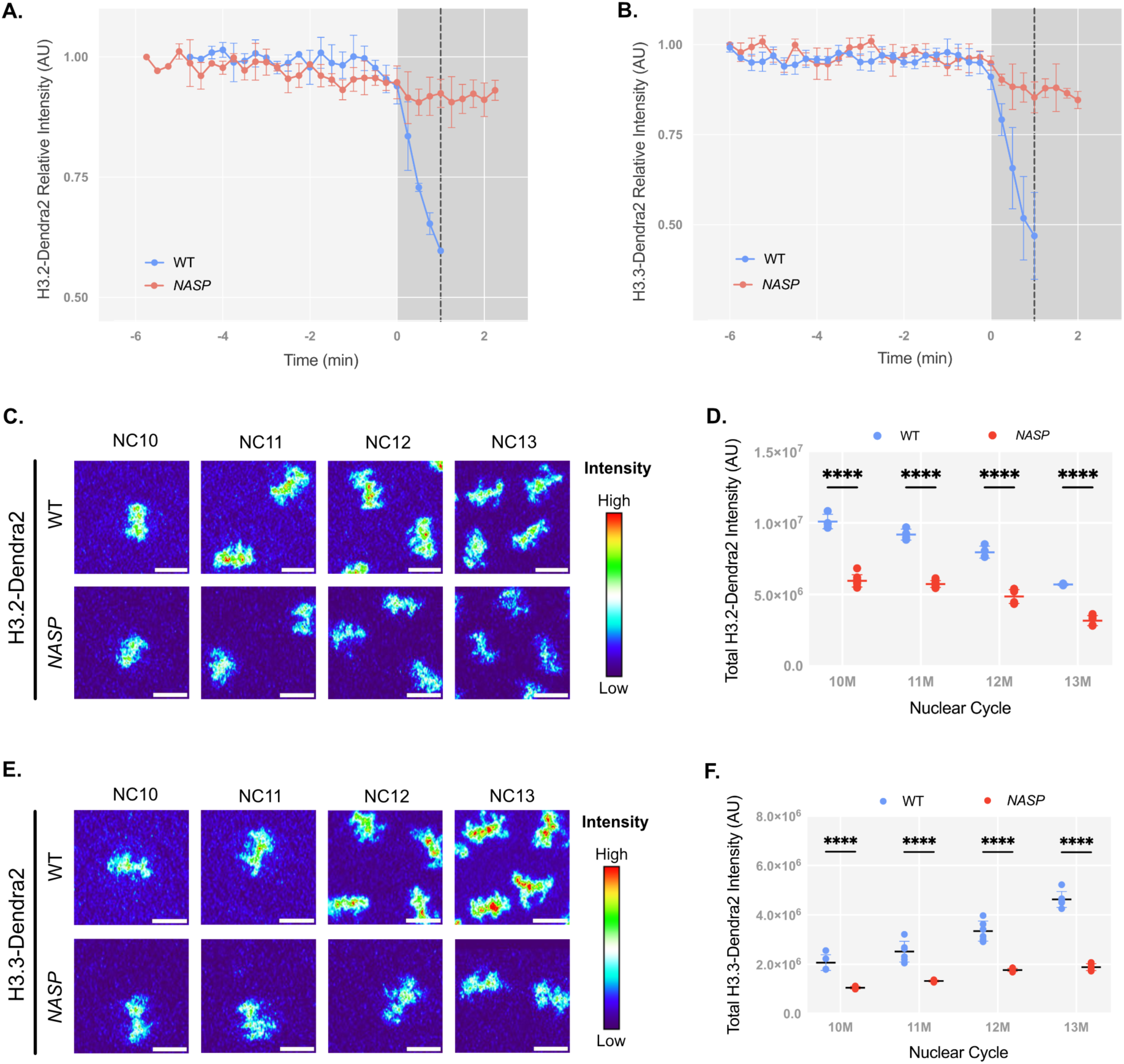
NASP deficient embryos have less nucleoplasmic and chromatin-associated H3. **(A)** Nuclear export measurement for H3.2-Dendra2 in WT and NASP-deficient embryos in NC12. Time point 0 represents Nuclear Envelop Breakdown (NEB). **(B)** Nuclear export measurement for H3.3-Dendra2 in WT and NASP-deficient embryos in NC12. **(C)** Maximum intensity projections of metaphase chromatin from WT (top) and NASP-deficient (bottom) embryos expressing H3.2-Dendra over NC10-13. Heat map created by pseudo-coloring images using nonlinear Thermal LUT on FIJI. **(D)** Total-intensities of H3.2-Dendra2 on mitotic chromatin of WT and NASP-deficient embryos for metaphase 10-13. **(E)** Maximum intensity projections of metaphase chromatin from WT (top) and NASP-deficient (bottom) embryos expressing H3.3-Dendra over NC10-13. Heat map created by pseudo-coloring images using nonlinear Thermal LUT on FIJI. **(F)** Total-intensities of H3.3-Dendra2 on mitotic chromatin of WT and NASP-deficient embryos for metaphase 10-13

Given the reduced import rate and nucleoplasmic concentration of H3.2 in embryos devoid of NASP, we wondered if this would lead to a reduction in the amount of chromatin-associated H3.2. To test this, we compared the total mitotic H3-Dendra2 intensities in embryos laid by wild-type and *NASP-*mutant mothers. We measured the absolute intensity of H3.2-Dendra2 in metaphase nuclei across nuclear cycles 10-13 (NC10-13) for these embryos (Figure 2C). The total H3.2-Dendra2 intensity across NC10-NC13 was significantly reduced in embryos laid by *NASP*-mutant mothers relative to embryos laid by wild-type mothers (Figure 2D). The reduced levels of chromatin bound H3.2 in embryos devoid of NASP is similar to the overall reduction in H3 in these embryos (Figure 2D) [40]. It is possible that the H3.3 could replace H3.2 in NASP-deficient embryos to maintain a constant nucleosome:DNA ratio. Therefore, we also measured chromatin bound H3.3 in embryos laid by wild-type and *NASP*-mutant mothers (Figure 2E). Similar to the overall reduction in H3.3 protein levels, we found that the H3.3 protein levels were also reduced in chromatin (Figure 2F). Thus, H3.3 does not compensate for reduced H3.2 levels in chromatin in NASP-deficient embryos. Despite the differences in chromatin-bound H3, embryos devoid of NASP have only a modest effect on cell cycle length starting in nuclear cycle 13 (Supplemental Figure 1I). We conclude that NASP indirectly affects the amount of H3.2 and H3.3 that gets incorporated into the chromatin during early nuclear divisions.

### H3 and NASP have different import kinetics

While H3 import kinetics are independent of NASP, it is possible that NASP and H3 form a complex in the cytoplasm that is imported into the nucleus. Biochemical data support the formation of a NASP-H3 complex in the cytoplasm, however, NASP dissociates from H3 upon the binding of Importin-4 or Importin-5 in mammalian cells [48]. Localization studies, coupled with in vitro assays, suggest that NASP is nuclear localized in mammalian cells and that NASP complexes with H3 in the nucleus [48, 49]. To date, there have been no studies that measure the localization and nuclear import kinetics of endogenous NASP and H3 to determine their nuclear import kinetics in vivo. Nor is it known if NASP localizes to nucleus during Drosophila embryogenesis. Therefore, we used CRISPR-based mutagenesis to add a Dendra2 tag the 3’ end of *NASP* at the endogenous locus, generating a C-terminally-tagged NASP-Dendra2 protein. Homozygous *NASP-Dendra2* flies were verified by sequencing and western blot (Supplemental Figure 2A-B). Homozygous *NASP-Dendra2* females were fertile indicating that the NASP-Dendra2 protein was functional. Live imaging of early embryos expressing NASP-Dendra2 revealed that NASP was largely nuclear localized, similar to NASP localization in Drosophila cultured cells and mammalian cells (Figure 3A; Supplemental Movie S1) [40, 49]. The NASP-Dendra2 intensity is nuclear during interphase and completely disperses upon nuclear envelope breakdown, while the H3.2-Dendra2 intensity persists throughout mitosis (Figure 3A). Thus, NASP is imported into the nucleus during early nuclear divisions and remains nucleoplasmic until it disperses upon nuclear envelope breakdown.

**Figure 3:**
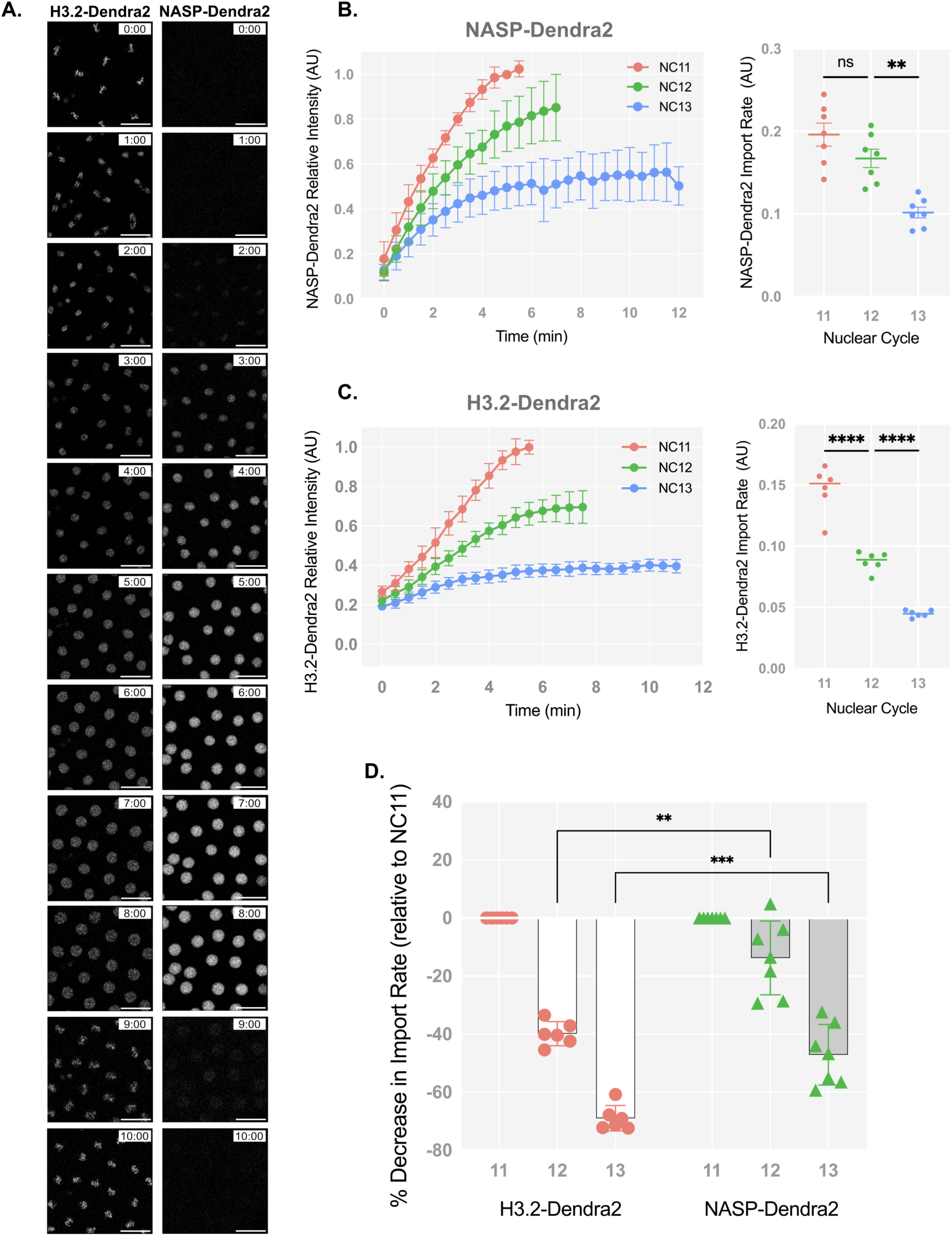
H3 and NASP have different nuclear import kinetics. **(A)** Time-lapse of H3.2-Dendra2 (left) and NASP-Dendra2 (right) from metaphase 12 to metaphase 13. **(B)** Relative intensity curves of NASP-Dendra2 for nuclear cycles NC 11-13 in WT embryos. **(C)** Relative intensity curves of H3.2-Dendra2 for nuclear cycles NC 11-13 in WT embryos (same as in Figure 1A). **(D)** The percent decrease in the nuclear import rate of H3.2-Dendra2 and NASP-Dendra2 for NC 11-13.

With NASP-Dendra2 we could now measure the nuclear import kinetics of NASP and H3 during the nuclear divisions of the early embryo. Absolute nuclear import rates are dependent on the concentration of the imported protein [57]. Given the concentration difference of Dendra2-tagged H3.2 and NASP in early embryos, we cannot directly compare the absolute nuclear import rates for these two proteins. We could, however, measure and compare the relative nuclear import rate of NASP-Dendra2 and H3.2-Dendra across nuclear cycles 11-13 (Figure 3B-C). We observed that the nuclear import dynamics of NASP-Dendra2 are starkly different from H3.2 (Figure 3D). Relative nuclear import rates of H3.2-Dendra2 reduce by half each nuclear cycle (NC11-13), consistent with exhaustion of the H3.2 pools as the embryo progresses through embryogenesis (Figure 1A; Figure 3C). The NASP-Dendra2 relative nuclear import rates, however, did not significantly change between nuclear cycle 11 and 12 (Figure 3B). Only in nuclear cycle 13 was there a decrease in NASP-Dendra2 import rate (Figure 3B and 3D). These kinetics were significantly different than those of H3.2-Dendra2 (Figure 3D). Subtracting the chromatin-bound values of H3.2 to measure the import rate of only the soluble fraction of H3.2 did not affect the import kinetics (Supplemental 2C). These observations show that H3.2 and NASP have different nuclear import kinetics during the nuclear divisions of the early embryo. Thus, in agreement with biochemical data, NASP and H3.2 are unlikely to be imported into the nucleus as a single complex and are rather imported independent of each other. Together with our data showing that the rate of H3 import or export is not dependent on NASP, we conclude that NASP has no direct effect on H3 import or export dynamics and that NASP is unlikely to act as a nuclear receptor for H3 during the nuclear divisions of the early embryo.

### NASP prevents H3 aggregation

We previously showed that H3 is not degraded in *NASP*-mutant stage 14 egg chambers [40]. Only upon fertilization and entry in embryogenesis is H3 degraded in NASP-deficient embryos [40]. To test if degradation of H3 is tied to egg activation or the onset of embryogenesis, we crossed wild-type and *NASP*-mutant mothers to *twine* mutant males to generate activated eggs that fail to progress into active embryos [58]. *twine* is essential for the completion of meiosis and *twine* mutant males fail to make sperm [58–60]. Overnight collections of eggs laid by wild-type females crossed to *twine* mutant males produce activated eggs that do not enter embryogenesis and thus do not have mitotic nuclei (Supplemental Figure 3A-B). We crossed wild-type or *NASP*-mutant mothers to *twine* mutant males and collected activated eggs. Western blot analysis of H3 levels revealed that H3 levels were reduced by 54% in activated eggs collected from *NASP*-mutant mothers (Figure 4A-B; Supplemental Figure 3C). This reduction was specific to H3 as H2B levels remain unaffected in eggs laid by *NASP*-mutant mothers (Figure 4A; Supplemental Figure 3C). Importantly, activated eggs do not contain mitotic nuclei yet NASP is required to prevent H3 degradation in these eggs. Therefore, NASP functions in the cytoplasm to protect H3 from degradation upon egg activation.

**Figure 4:**
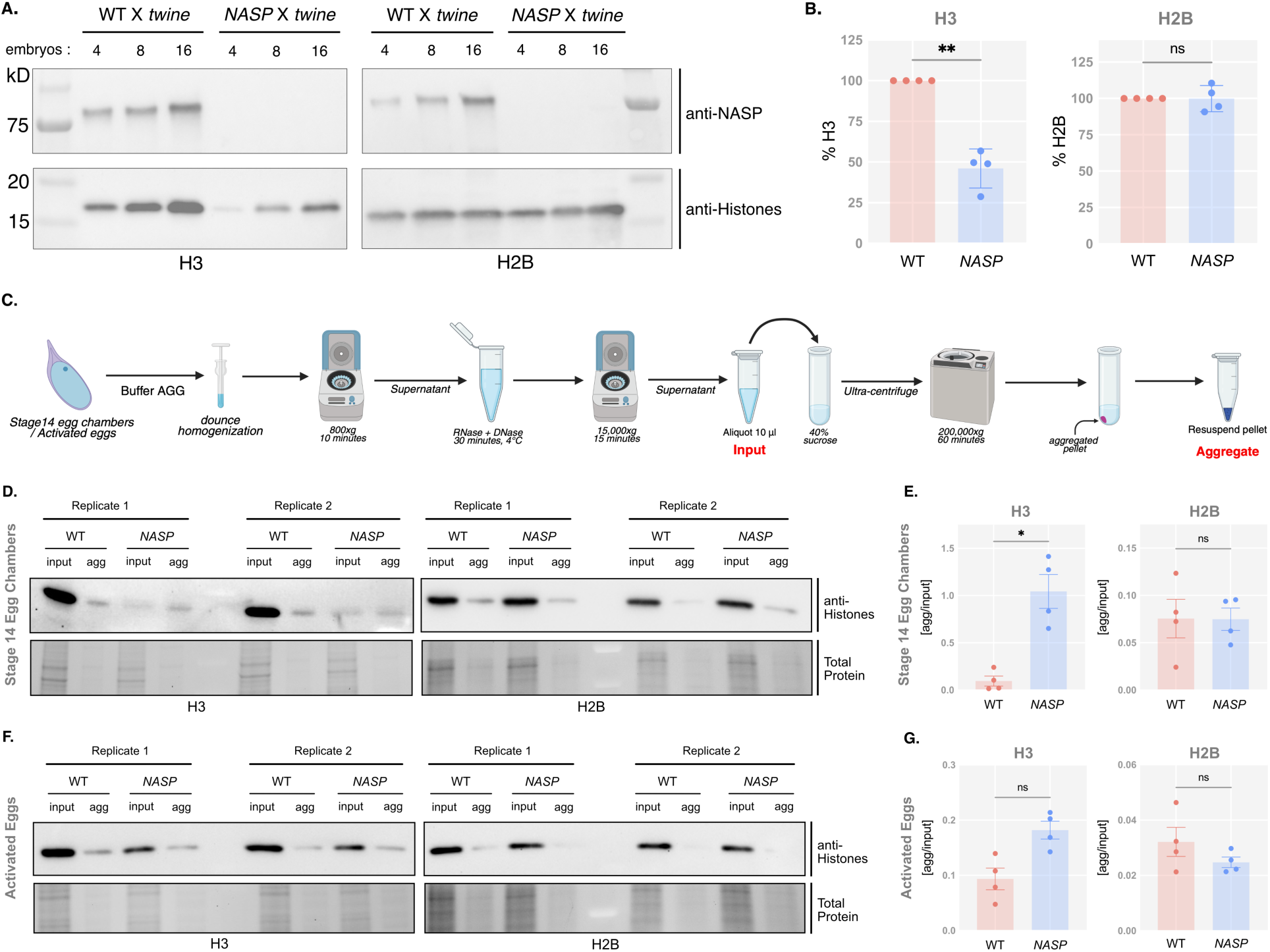
NASP functions in the cytoplasm. **(A)** Western blot analysis of activated eggs collected from *WT* or *NASP-*mutant females crossed to *twine* mutant males. **(B)** Quantification of H3 and H2B protein levels in activated eggs collected from *WT* or *NASP-*mutant females crossed to *twine* mutant males. Protein levels are normalized to the WT intensities. **(C)** Schematic of the aggregate isolation protocol used to separate aggregates from stage 14 egg chambers and early embryos. **(D)** Western Blot analysis for input and aggregate fractions from stage 14 egg chambers from *WT* and *NASP* mutant female flies. **(E)** Ratios of H3 and H2B in the aggregate fraction relative to input fractions in stage 14 egg chambers of WT and *NASP* mutant female flies. **(F)** Western blot analysis for input and aggregate fractions derived from activated eggs collected from *WT* or *NASP-*mutant females crossed to *twine* mutant males. **(G)** Ratios of H3 and H2B in the aggregate fraction relative to input fractions from activated eggs collected from *WT* or *NASP-*mutant females crossed to *twine* mutant males.

Histones are highly prone to aggregation in vitro [61]. Furthermore, in *NASP* mutant stage 14 egg chambers, there is a higher fraction of insoluble H3 as compared to the wild-type egg chambers [40]. Therefore, the primary function of NASP during oogenesis could be to prevent H3 aggregation. To test this possibility, we adopted an aggregate isolation assay used to isolate protein aggregates from metazoan tissue lysate (Figure 4B) [62]. By using differential centrifugation, this protocol separates aggregates from larger complexes including organelles, ribonucleoproteins and condensates [63–69]. Stage 14 egg chambers were collected from wild type and *NASP*-mutant females, lysed and subject to differential centrifugation to obtain a lysate for ultracentrifugation. These lysates were subject to ultracentrifugation to isolate protein aggregates (Figure 4B). Western blot analysis of the input and aggregate fractions revealed that ∼9.5% of H3 was in the aggregation fraction in wild type egg chambers (Figure 4D-E; Supplemental Figure 4A). The amount of H3 in aggregate fraction *NASP*-mutant egg chambers, however, was increased ∼11-fold relative to wild type (Figure 4D-E). Aggregation was specific to H3 as there was no significant enrichment of H2B in aggregates derived from *NASP-*mutant egg chambers. Given that H3 degradation in the absence of NASP is tied to egg activation (Figure 4A), we next asked if the aggregate fraction of H3 was preferentially targeted for degradation in the absence of NASP. We collected activated eggs from wild-type and *NASP*-mutant mothers that were crossed to *twine* mutant males and performed the same aggregate isolation protocol. This analysis revealed no significant difference in the amount of H3 in the aggregate fraction when comparing lysates derived from activated eggs that to do or do not contain NASP (Figure 4F-G; Supplemental Figure 4B). Thus, we conclude that the upon egg activation, protein aggregates that contain H3 are preferentially targeted for degradation.

Taken together, we conclude that NASP does not directly protect H3 from degradation. Rather, NASP prevents H3 from aggregation. In the context of oogenesis and embryogenesis, aggregation and degradation are developmentally separated allowing the two processes to be distinguished. In the absence of NASP, aggregation first occurs during oogenesis when H3 is being deposited into the egg chamber. Only upon egg activation are H3 protein aggregates targeted for degradation.

## DISCUSSION

NASP is a H3-specific chaperone and the interaction between NASP and H3 has been well characterized on a biochemical and structural level [42–47]. How NASP functions in vivo to control H3 dynamics, however, is less well understood. By utilizing the early Drosophila embryo as a model system to study H3 dynamics, we found that NASP does not directly affect the nuclear import or export rates of histone H3.2 or H3.3. Rather, in the absence of NASP H3 protein levels are reduced, which leads to an indirect reduction in H3 nuclear import rate, less soluble H3 in the nucleoplasm and reduced H3 in chromatin. While NASP does localize to the nucleus in Drosophila embryos, NASP primarily functions in the cytoplasm to chaperone H3. We found that H3 aggregation and degradation are developmentally uncoupled during Drosophila oogenesis and embryogenesis. Using this powerful developmental system, we have provided evidence that the key function of NASP in vivo is to prevent H3 aggregation rather than directly preventing H3 degradation.

To sustain the rapid nuclear cycles of early embryogenesis, embryos must be stockpiled with excess histones [20]. This provides a unique opportunity to understand how H3 nuclear import kinetics are affected by NASP. To our surprise, NASP does not directly affect the nuclear import kinetics of H3. Likewise, NASP does not act as receptor to sequester H3 in the nucleus and prevent H3 from export. NASP directly binds to H3 and a number of NASP:H3 complexes have been identified from cell extracts [42, 47–48]. H3 binding to Importin-4 or Importin-5, however, is mutually exclusive with NASP binding to H3 [42, 48]. Therefore, it is not clear how NASP could directly facilitate nuclear import of H3 if NASP is not part of the protein complex that is directly imported through the nuclear pore. NASP is localized to the nucleus in somatic cells and the nuclear-specific function of NASP has yet to be defined. Given that H3 is not exported from the nuclei in Drosophila embryos [31] it was possible that by binding to H3, NASP prevented nuclear export of H3 [49]. Our results show that even in the absence of NASP, H3 is not exported from the nucleus. Thus, understanding the function of NASP in the nucleus remains enigmatic. In cancer cells, NASP functions in the nucleus to capture evicted nucleosomes upon PARP or topoisomerase inhibition [70]. This function is specific to cancer cells, thus the function of NASP in unperturbed human cells is still undefined.

In mammalian somatic cells, NASP prevents the degradation of H3, which occurs through chaperone-mediated autophagy [50]. It is still unclear how NASP physically protects H3 from degradation. In this work, we demonstrated that H3 aggregation and H3 degradation are developmentally separated. Our work reveals that in the absence of NASP, H3 aggregation precedes degradation. Thus, it appears that at least one critical function of NASP in vivo is to prevent H3 aggregation. While we think H3 aggregation likely proceeds degradation in somatic cells depleted of NASP, the limiting amounts of soluble histones make this difficult to address. By using activated eggs that are devoid of mitotic nuclei, we showed that NASP functions in the cytoplasm. In somatic cells and the early Drosophila embryo, however, NASP localizes to the nucleus. Because the cytoplasmic fraction of NASP is diffused in the Drosophila embryo, we cannot easily quantify the fraction of NASP in the nucleus during each nuclear cycle. Regardless, we propose that NASP functions both in the nucleus and the cytoplasm to prevent H3 aggregation. In the cytoplasm, protein aggregates are largely degraded through autophagy [71–73]. Consistent with this, chaperone-mediated autophagy is the main H3 degradation pathway in somatic cells depleted of NASP [50]. Factors that drive autophagy are not present within the nucleus [74]. Thus, NASP function in the nucleus could be extremely important to prevent potentially toxic nuclear aggregates of H3 that cannot be cleared through autophagy until nuclear envelope breakdown. Aggregates of H3 in the nucleus are likely to have similar properties to chromatin and could be more challenging to biochemically isolate and characterize. The pathway that degrades maternally inherited H3 aggregates in Drosophila has yet to be defined and Drosophila lacks LAMP-2A, a critical factor for chaperone mediated autophagy [75].

Only about ∼30% of embryos laid by NASP-mutant mothers hatch and we have yet to define the molecular mechanism(s) that prevent these embryos from proper development [40]. While embryos devoid of NASP have ∼50% of the amount of H3 as embryos laid by wild-type embryos, we do not believe histone deficiency underlies the problems during embryogenesis. First, nearly half of the embryos laid by NASP mutant mothers are arrested with one or two nuclei [40]. Depleting half of the histone H3 pool in the early embryo should still provide enough histone to progress until nuclear cycle 13 [31]. Second, embryos laid by Jabba-mutant mothers (the H2A/H2B-specific chaperone) have vastly depleted maternal pools of H2A and H2B and yet proceed through embryogenesis relatively normally [39]. Third, embryos actively translate maternally deposited histone RNA and this can compensate for reduced histone pools [76]. One intriguing possibility is that the maternally transferred H3 protein aggregates cause toxicity in the early embryo. Protein aggregates are known to be toxic in several contexts [77–80]. Furthermore, aggregates containing high levels of histones could mimic chromatin and recruit histone chaperones (e.g. HIRA) or other histone modifying enzymes into the aggregate where they would be degraded or sequestered. Recent work in the mouse embryo has revealed that protein aggregates in the mouse oocyte can inhibit embryo survival if not degraded [81].

Embryos laid by *NASP*-deficient females have significantly reduced H3 levels that indirectly affects the import rate and nucleoplasmic supply of H3. Remarkably, NASP-deficient embryos also have an ∼50% reduction in chromatin-associated H3. Although the embryos laid by NASP deficient mothers have less H3 in their chromatin, the nuclei divide relatively normally, albeit a little slowly. The reduced incorporation of H3 into chromatin in these embryos presents an interesting question about genome packaging in these embryos. For example, are nucleosomes randomly distributed or are there hotspots where nucleosomes must be positioned? How does a reduced number of nucleosomes affect processes like zygotic transcription and DNA replication? Using a tractable developmental model system will allow us to address these questions while discerning the function of NASP during embryogenesis and beyond.

## MATERIALS AND METHODS

### Strain List and stock generation

Wild type – Oregon R (OrR)

*NASP* null mutant (*NASP)* - *w[1118]; Df(3R)Exel6150, P{w[+mC] = XP-U}*

*Exel6150/TM6B, Tb[1]/NASP^2^*

Df(3R) *- w[1118]; Df(3R)Exel6150, P{w[+mC] = XP-U} Exel6150/TM6B, Tb[1]*

NASP-Dendra2 *- y[1] M{vas-Cas9}ZH2A w[1118]; NASP-Dendra2*

H3.2-Dendra2 *- y,w; 1xHisC.H3-Dendra2*

H3.3-Dendra2 - *y,w; H3.3A-Dendra2/CyO*

*twine – twe[1] cn[1] bw[1]/CyO* and *y[1] w[67c23]; P{w[+mC]=lacW} twe[k08310]/CyO*

To generate a Dendra2-tagged allele of *NASP*, a single gRNA targeting C-terminal of *NASP* was cloned into pU6-BbsI plasmid as described [82]. The gRNA was identified using the DRSC Find CRISPRs tool (http://www.flyrnai.org/crispr2/index.html). The gRNA-expressing plasmid was injected into a *vas-Cas9* expression stock (Best Gene Inc.) and balanced over *TM3*. The flies were self-crossed and screened for lack of *TM3* phenotype to establish a homozygous stock.

### Embryo Collection

*NASP/Df(3R)* or OrR female flies were yeast fed for 4 days at room temperature. The ovaries were dissected in 1X PBS and stage 14 egg chambers were isolated. For embryo collection, the fly strains or crosses were set up at room temperature in bottles capped by a grape juice agar plate with some wet yeast. For western blotting, 0–2-hour (AEL – after egg-laying) embryos were collected. To obtain activated eggs from *twine* mutant crosses, 0–1-hour (AEL) embryos were collected. To image activated eggs, overnight embryos (16-24 hr AEL) were collected. All the embryo collection samples were dechorionated with 50% bleach for up to 2 minutes and thoroughly washed with water twice. The dechorionated embryos were either flash frozen with liquid nitrogen and stored in -80°C or processed for further experiments.

### Live imaging

#### Images for Nuclear Import and Chromatin Analysis

For live imaging, Drosophila embryos were collected from fresh grape juice plates after allowing flies to lay for 2 hours. Embryos were then dechorionated with 30% bleach solution for 1 minute and 30 seconds and washed with water. Embryos were mounted on a glass-bottom dish in deionized water and images were acquired using a Zeiss LSM 980 confocal microscope with Airyscan-2 the 40x, 1.2 NA water objective. Embryos laid by *H3.2-Dendra2, H3.2-Dendra2; NASP;* and *NASP-Dendra2* mothers were imaged with the 488nm laser at 2% power using the CO8-Y Airyscan Setting. *H3.3-Dendra2* embryos were imaged at 0.5% laser power. All z-stacks comprised of 16 planes spaced 1µm apart. The stage was heated at 25°C.

For nuclear import and chromatin analysis, all were imaged every 45 seconds for 2 hours, with a 712 x 712-pixel resolution, at 3x zoom, and a 35.88ms frame time. The pixel size of all images was 0.099 µs x 0.099µs

#### Images for Nuclear Export

For nuclear export, imaging was performed as described for nuclear import with the following exceptions. Nuclear export imaging was done with interactive bleaching mode on ZEN Blue Software. Dendra2 tag was photoconverted from green to red using 60 iterations of the 405nm laser at 3% power with an exposure speed of 1.37µs/pixel. A 4µm diameter circle stencil was used to outline the nuclei meant to be photoconverted. The nucleus was converted in the middle of the nuclear cycle and imaged for 15-second intervals until the end of the nuclear cycle. Images were captured at a 576 x 576 pixel resolution, at 4 x zoom, with a 66.55ms frame time, and a pixel size of 0.092 µm x 0.092 µm.

#### Photobleaching

To determine potential photobleaching during image acquisition, two embryos of the same age were imaged from the interphase nucleus and metaphase chromatin for NC10. Then a sub-region of one embryo was imaged with the experimental import settings until NC13. Once imaged embryo reached NC13 both were imaged at the interphase nucleus and metaphase chromatin of NC13. The continuously imaged area was compared to the outside area and the unimaged embryo. These comparisons showed minimal photobleaching, and therefore, no numerical photobleaching corrections were applied to our data.

#### Segmentation and Image Analysis

Images were processed using ZEN 3.3 (blue edition), using 3D Airyscan Processed with a strength of 3.7. Scenes were separated and converted to individual TIFF files.

The z-stack for the time point of metaphase chromatin for each nuclear cycle was sum projected in FIJI. The chromatin and cytoplasm of these images were then segmented using the pixel classification + object classification feature on Ilastik. A CSV file was exported containing the total intensity.

To obtain nuclear import curves live images were divided into individual cycles and segmented using pixel classification + object classification on Ilastik. CSV files contained total intensity values which were subtracted by the nuclear intensity of the first frame of each cycle to plot the total intensities starting at 0. To obtain relative intensities, the total intensity values were normalized to the end of nuclear cycle 11 for each embryo. Cell Cycle duration was measured from metaphase to metaphase of the next cycle.

#### Statistical analysis

To calculate import rates the initial slope of the import curves was calculated using a simple linear regression. Two-way ANOVA tests were performed to evaluate the statistical significance between different genotypes.

### Western Blotting

Protein lysates were prepared by homogenizing dechorionated embryos in 1.5ml Eppendorf tube using a pestle in 2X Laemmli Sample Buffer (Bio-Rad, Cat#1610737) supplemented with 50mM DTT. Lysates were boiled for 10 minutes and loaded on a 4-15% Mini-PROTEAN TGX Stain-Free Gel (Bio-Rad, Cat#4568086). After electrophoresis, gels were activated and imaged using a BioRad ChemiDoc MP Imaging System using default manufacturer parameters. Protein was transferred to Immobilon-P PVDF Transfer Membrane (Millipore, Ref#IPVH00010) using the BioRad Trans-Blot Turbo Transfer System and membranes were imaged using the BioRad ChemiDoc MP Imaging System using default manufacturer parameters. Transfer membrane was blocked with 5% non-fat milk in 1X TBS-T (140 mM NaCl, 2.5 mM KCl, 50 mM Tris HCl pH 7.4, 0.1% Tween-20) for 10 minutes. Blots were incubated with primary antibodies (anti-NASP – 1:2000, anti-H3-HRP – 1:1000, anti-H2B – 1:1000) for one hour at room temperature, washed and incubated with secondary antibodies (HRP anti-mouse – 1:20,000, HRP anti-rabbit – 1:25,000) for 30 minutes at room temperature. Blots were then washed with TBS-T and incubated with Clarity ECL Western Solution (BioRad, Cat#1705061) for four minutes prior to imaging. Blots were imaged for using the BioRad ChemiDoc MP Imaging System.

### Aggregate Isolation

A standard aggregate isolation protocol was adapted from Chen *et al* [61] to isolate protein aggregates from Drosophila egg chambers and activated eggs. Stage 14 egg chambers or activated eggs were homogenized using a dounce homogenizer with a B-type pestle in 100μL of buffer AGG (30 mM Tris-Cl pH 7.5, 1 mM DTT, 40 mM NaCl, 3 mM CaCl_2_, 3 mM MgCl_2_, 5% glycerol, 1% Triton X-100 supplemented with Roche cOmplete EDTA-free Protease Inhibitor Cocktail, Cat#04693132001). The homogenized lysate was transferred to a 1.5mL Eppendorf tube and centrifuged at 800x*g* for 10 minutes at 4°C to remove cell debris. The supernatant was transferred to a new 1.5mL Eppendorf tube and treated with 100μg/mL RNase A (Thermo Scientific, Cat#EN0531) and 100μg/mL DNase I (NEB, Cat#M030S)for 30 min on ice. Nuclease treated samples were centrifuged at 10,000x*g* for 15 minutes at 4°C (same Beckman Coulter FA-241.5 rotor). The resulting supernatant is the “input lysate” fraction and an aliquot of this fraction was saved for western blotting. The remaining lysate was loaded on top of 1ml 40% sucrose pad and an additional 750μL of Buffer AGG was added to the lysate in the ultracentrifugation tube (Beckman Coulter Polypropylene Centrifuge Tubes, Cat#347357). Samples were subjected to ultra-centrifugation for one hour at 200,000x*g* at 4°C (Optima TL Ultracentrifuge using a Beckman TLS-55 rotor). Most of the supernatant was removed leaving 10-15μL of sample at the bottom of the ultracentrifuge tube. An additional 20μL of Buffer AGG was added to rigorously resuspend the remaining fraction which constitutes the “aggregate” fraction.

## DATA AVAILABILITY

All raw and processed images are available upon request

## ACKNOWLEDGMENTS

Essential fly stocks were provided by the Bloomington Drosophila Stock Center (NIHP40OD018537). This research was supported by NIH grants 2R35GM128650 (to J.T.N) and R35GM150853 to A.A.A

## AUTHOR CONTRIBUTION

Conceptualization, MD and JTN; Investigation, MD, EC-C; Analysis, MD, EC-C, ADB; Methodology, RT; Writing – original draft, MD and JTN; Writing – review & editing MD, EC-C, ADB, RT, AAA, JTN; Funding acquisition, JTN and AA; Supervision, JTN and AAA

## CONFLICT OF INTEREST

The authors declare no competing interests

## Notes

### Competing Interest Statement

The authors have declared no competing interest.

