## Supplemental_figures for "NASP functions in the cytoplasm to prevent histone H3 aggregation during early embryogenesis"

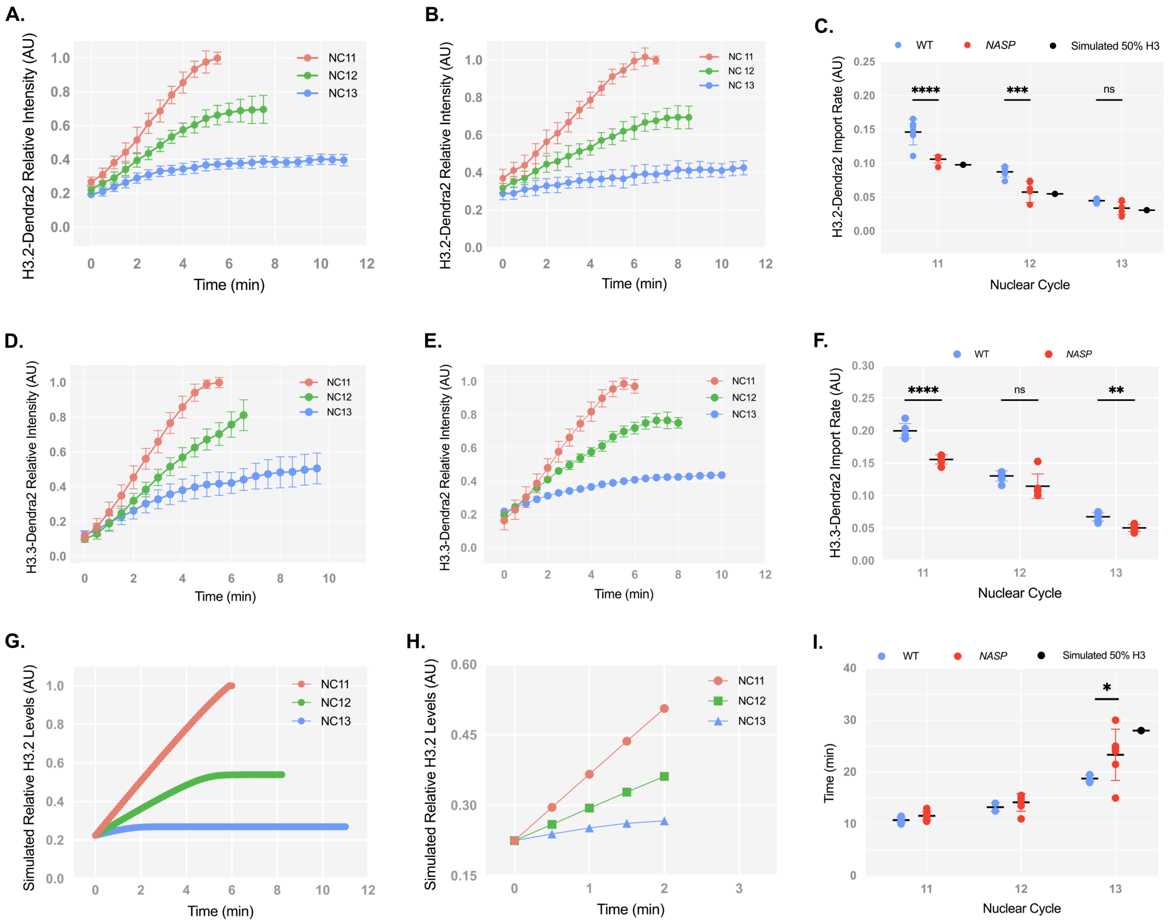


**Supplemental Figure 1: Normalized nuclear import rates for H3.2 and H3.3 Dendra**

**(A)** Normalized relative intensity curves of H3.2-Dendra2 for nuclear cycles (NC) 11-13 in WT embryos. **(B)** Normalized relative intensity curves of H3.2-Dendra2 for nuclear cycles (NC) 11-13 in NASP-deficient embryos.**(C)** Normalized nuclear import rates of H3.2-Dendra2 in WT (blue) and NASP-deficient (red) embryos. Simulated import rate for H3.2 at 50% wild-type levels (black). **(D)** Normalized relative intensity curves of H3.3-Dendra2 for nuclear cycles (NC) 11-13 in WT embryos. **(E)** Normalized relative intensity curves of H3.3-Dendra2 for nuclear cycles (NC) 11-13 in NASP-deficient embryos. **(F)** Normalized nuclear import rates of H3.3-Dendra2 in WT (blue) and NASP-deficient (red) embryos. **(G)** Normalized relative intensity curves for simulation of H3 nuclear import at 50% initial concentration. **(H)** Slope for nuclear import rate calculation for simulation of H3 nuclear import at 50% initial concentration. **(I)** Cell cycle duration for NC11-13 in WT (blue) and NASP-deficient (red) embryos. Simulated cell cycle duration for NC13 at 50% wild-type H3.2 levels (black).

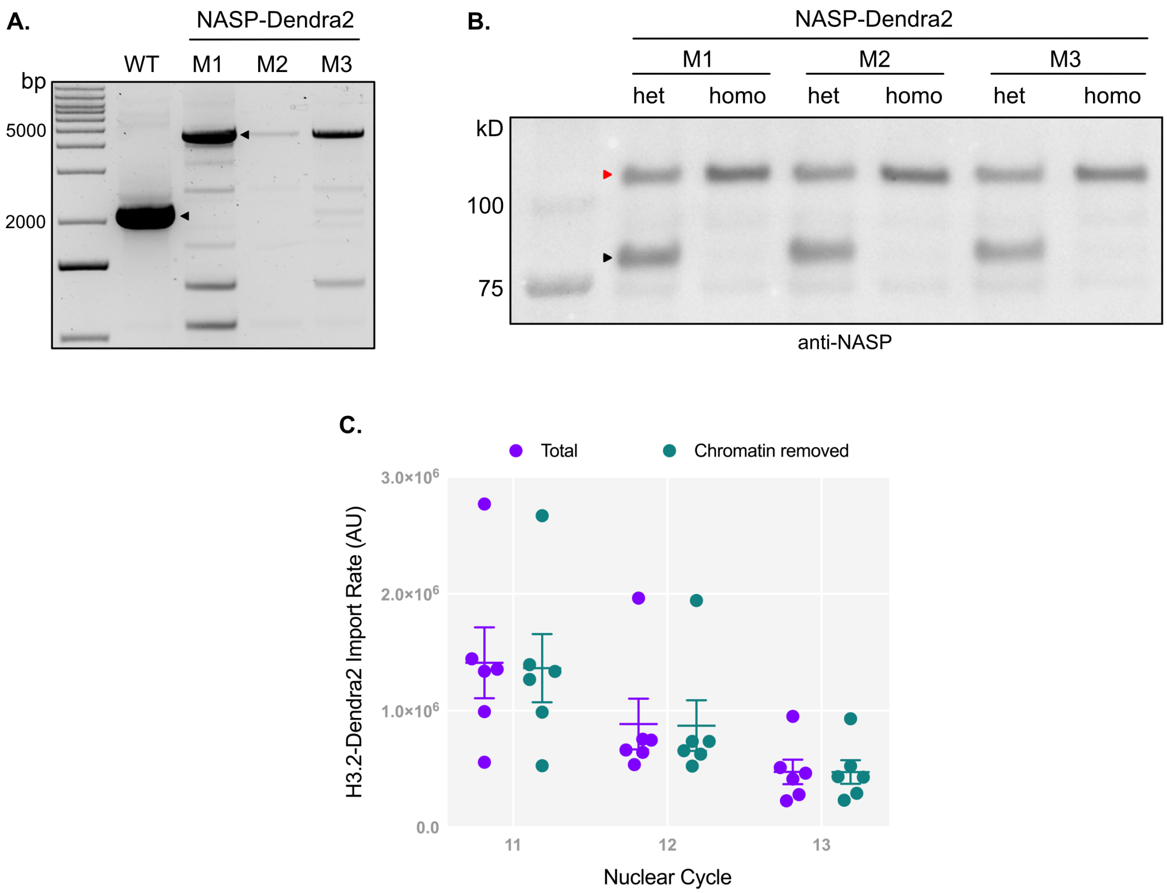


**Supplemental Figure 2: Validation of NASP-Dendra fly line**

**(A)** PCR analysis of the NASP endogenous locus shows ~3kb shift in the NASP-Dendra tagged flies as compared to the WT flies. WT -wild type, M1 – male #1, M2 – male #2, M3 – male #3 **(B)** Western blot analysis of ovary extracts from heterozygous (het) and homozygous (homo) *NASP-Dendra* female flies confirm the expression of NASP-Dendra (red arrowhead represents Dendra-tagged NASP and the black arrowhead represents wild-type NASP) **(C)** H3.2 Dendra nuclear import rates calculated from unnormalized total nuclear intensities (purple) and non-chromatin associated intensities (teal).

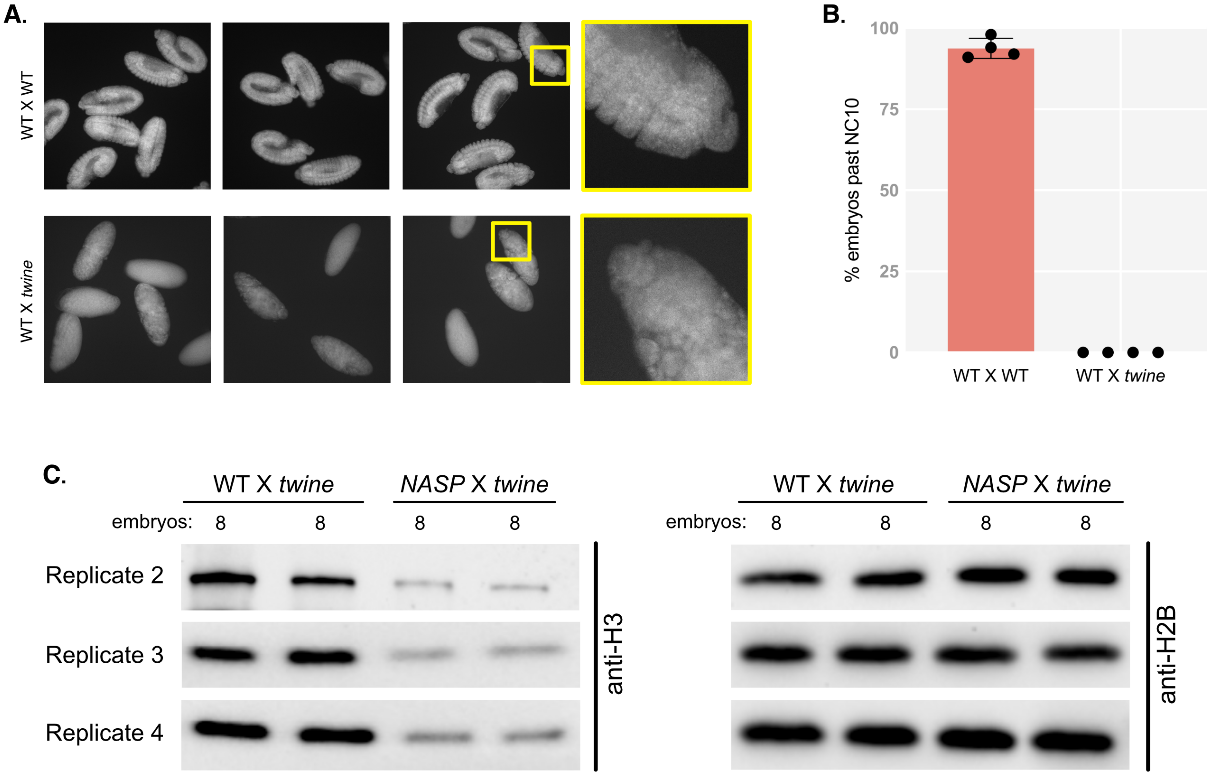


**Supplemental Figure 3: Females crossed to *twine* mutant males fail to develop**

**(A)** DAPI stained eggs from wild-type females crossed to wild-type males (WT X WT) and wild type females crossed to *twine* mutant males (WT X *twine*). Imaging was at 10X or 60X(yellow boxes) **(B)** Percentage of eggs laid that progress past nuclear cycle 10 from *WT* X *WT* and *WT* X *twine* mutant crosses. **(C)** Western blot analysis of activated eggs collected from *WT* or *NASP*-mutant females crossed to *twine* mutant males used in quantification in Figure 4B (Replicates 3 and 4).

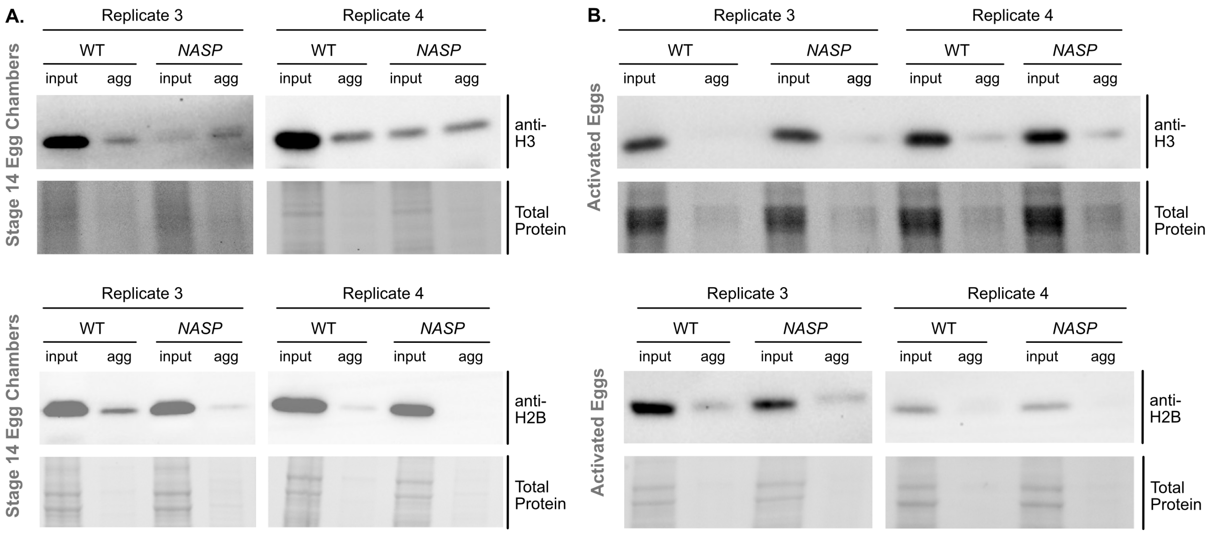


**Supplemental Figure 4: Western blots of histone aggregation across multiple replicates**

**(A)** Western blot analysis for input and aggregate fractions from stage 14 egg chambers from *WT* and *NASP* mutant female flies used in quantification in Figure 4E (Replicates 3 and 4). **(B)** Western blot analysis for input and aggregate fractions derived from activated eggs collected from *WT* or *NASP*-mutant females crossed to *twine* mutant males used in quantification in Figure 4G. (Replicates 3 and 4)
